# Structure and antigenicity of the 2026 Bundibugyo virus glycoprotein

**DOI:** 10.64898/2026.10.22.761020

**Authors:** Amin Addetia, Keiya Uriu, Cameron Stewart, Kaitlin R Sprouse, Yu Kaku, Mohammad Golam Kibria, Olivia Putri, S M Tanjil Shah, Yuri V Fukushima, Shusuke Kawakubo, Young-Jun Park, The Genotype to Phenotype Japan (G2P-Japan) Consortium, Akatsuki Saito, Kei Sato, David Veesler

## Abstract

An outbreak of ebolavirus disease caused by a new zoonotic spillover of Bundibugyo virus (BDBV) is currently spreading at an unprecedented pace in the Democratic Republic of the Congo and Uganda. The outbreak has resulted in 7,733 cases with a case-fatality ratio of 48%, yet no BDBV-specific vaccines or therapeutics are available. The ebolavirus glycoprotein (GP) promotes viral entry and is the target of neutralizing antibodies and vaccine development. Here, we engineered a soluble, prefusion-stabilized GP trimer from a contemporary BDBV isolate and determined its cryo-electron microscopy structure bound to the broadly neutralizing antibody EBOV-515. It reveals the molecular basis for EBOV-515 cross-reactivity and suggests that the antibody inhibits viral entry by interfering with NPC1 engagement in addition to trapping GP in the prefusion conformation and hindering glycan cap cleavage. Mapping lineage-defining substitutions identified changes in GP1 and GP2, including R506S within an antigenic site targeted by multiple broadly neutralizing antibodies recognizing the fusion loop. Nevertheless, we found that several glycan cap-, GP base-, and HR2-directed antibodies cross-reacted and neutralized BDBV DRC/2026 GP pseudovirus with potency comparable or greater to that observed against earlier BDBV isolates and the Zaire ebolavirus Makona/2024 isolate. These findings define the antigenic properties of contemporary BDBV GP and delineate conserved epitopes as ideal targets for antibody therapeutics and for the development of pan-ebolavirus vaccines.

---

Filoviruses are highly lethal pathogens causing hemorrhagic fever and organ failure with fatality rates ranging from 25 to 90%^1^. They are divided in two genera, designated Ebolavirus and Marburgvirus, and typically spillover to humans from bats^2–4^. The Ebolavirus genus comprises six viral species, namely Zaire (ZEBV), Sudan (SUDV), Bundibugyo (BDBV), Tai Forest (TAFV), Reston (RESTV), and Bombali (BOMV) with the first three species causing fatal disease in humans. The 2013-2016 EBOV outbreak in West Africa is the largest documented outbreak thus far with 28,616 infections and 11,310 deaths.

An outbreak of ebolavirus disease caused by BDBV is currently ongoing in the Democratic Republic of the Congo and in Uganda with cases exported to Germany and France. Although this outbreak was only recognized by the WHO in May, genomic sequencing indicates that the virus has been circulating since February. To date, the ongoing outbreak has led to greater than 7,279 confirmed cases and 3,512 deaths **(Fig 1a)**. It is the second-largest outbreak in history and is currently spreading more rapidly than the 2013-2016 EBOV outbreak in West Africa. It is therefore predicted to become the largest outbreak of ebolavirus disease ever documented which is particularly concerning as no BDBV-specific or pan-ebolavirus vaccines or treatments are approved. As a result, the WHO declared this outbreak a public health emergency of international concern.

**Figure 1.**
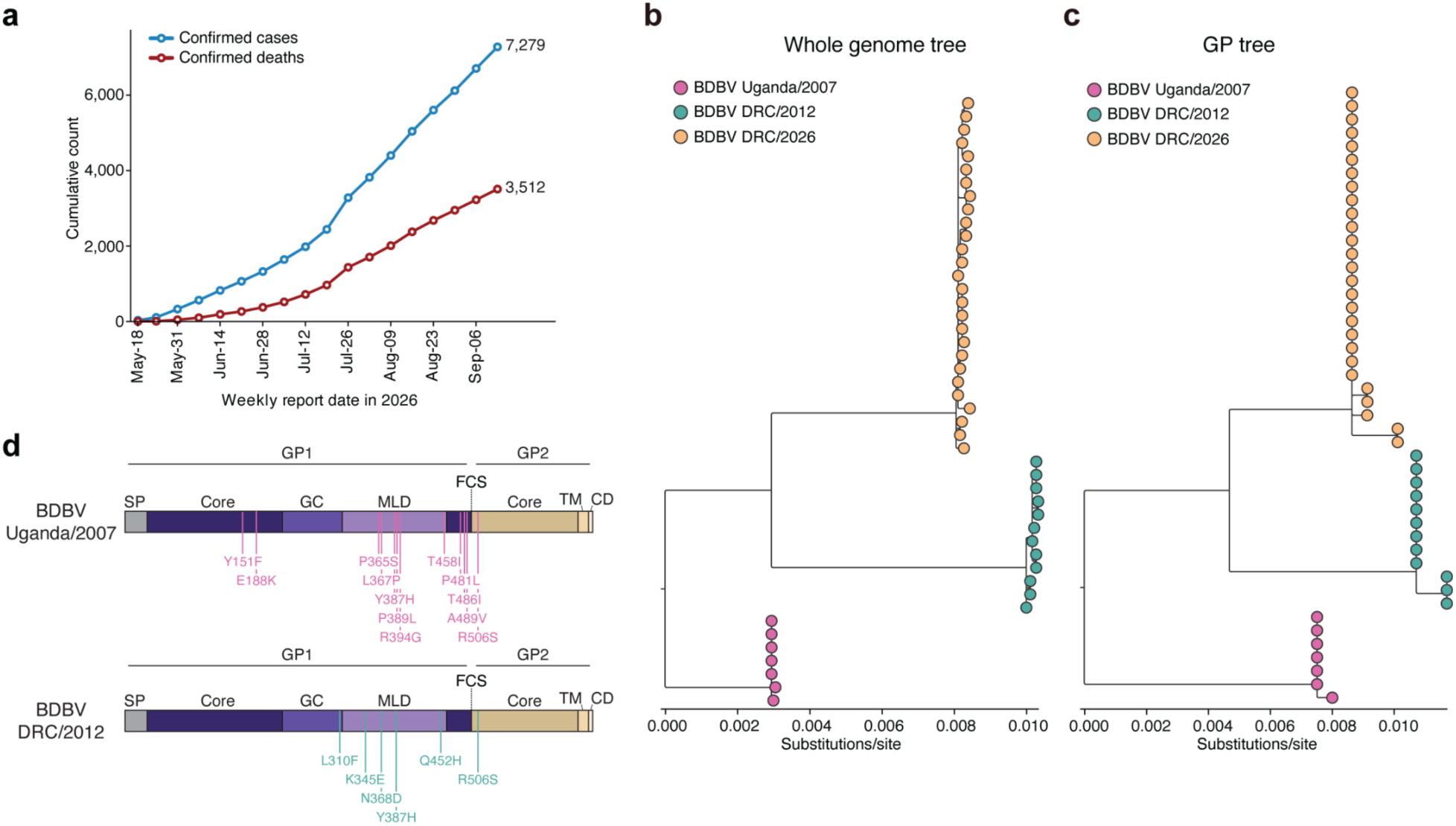
Epidemiology and genetic characterization of 2026 BDBV. **a**, Cumulative numbers of confirmed cases and deaths of BDBV disease. Data were obtained from WHO AFRO Weekly External Situation Reports Reports 01–18 (data as of 2026-Sep-13). **b,** Maximum-likelihood phylogenetic tree based on whole genome nucleotide sequences of BDBV. **c,** Phylogenetic tree based on GP coding region nucleotide sequences of BDBV. **d,** Schematic of BDBV GP domain organization and mutations found in the BDBV DRC/2026 GP sequence relative to GP sequences from the prior outbreaks of BDBV-induced disease (BDBV Uganda/2007 and BDBV DRC/2012). CD, cytoplasmic domain; FCS, furin cleavage site; GC, glycan cap; MLD, mucin-like domain; TM, transmembrane domain. SP, signal peptide.

Filoviruses enter cells using glycoprotein (GP) homotrimers, which mediate host receptor engagement and fusion of the endosomal and viral membranes to initiate infection^5–10^. GP is cleaved by furin during biogenesis, yielding GP1 and GP2 that remain covalently linked via a disulfide bond at the base of the prefusion GP trimer. Upon viral internalization, cathepsin-mediated GP cleavage enables shedding of the glycan cap and mucin-like domain, enabling engagement of the NPC1 host receptor before further cleavage promotes the large-scale conformational changes leading to membrane fusion^11–13^. GP is the target of neutralizing antibodies which correlate with protection in small animal and non-human primate challenge models^14–26^.

A live-attenuated vaccine based on a recombinant vesicular stomatitis virus (VSV) encoding EBOV GP was developed and licensed under the commercial name Ervebo^27–29^. Ebolavirus species exhibit considerable evolutionary divergence with GPs from two species typically sharing only 55-65% amino acid sequence identity. As a result, GP-elicited antibody responses against vaccine-mismatched GP species are reduced and vaccine-induced protection is dampened in animal models, relative to vaccine-matched immunity ^30–33^. A clinical trial run during the 2018 EBOV outbreak in the Democratic Republic of the Congo showed that the mAb114 monotherapy and the Inmazeb (REGN-EB3) cocktail therapy outperformed the previously developed Zmapp cocktail to reduce EBOV-induced mortality and they are now both approved by the FDA for treatment of EBOV infections^34,35^. However, these licensed mAbs are specific for EBOV GP and not for use against other ebolavirus species.

## Results

### Design of a prefusion-stabilized BVDV GP ectodomain trimer

Analysis of genome and GP sequences obtained from individuals infected during the ongoing outbreak indicate that BDBV DRC/2026 isolates form a distinct phylogenetic cluster from BDBV isolates associated with the 2007 and 2012 outbreaks, consistent with a new zoonotic spillover event^36^ **(Fig 1b-d)**. To characterize the GP architecture of the circulating BDBV strain and map lineage-defining mutations, we sought to recombinantly produce a soluble, prefusion-stabilized ectodomain GP trimer. We first evaluated a construct encoding for a mucin domain-deleted GP, obtained from a genome sequence collected on May 7th 2026 in the Democratic Republic of the Congo (https://pathoplexus.org/seq/PP_006XHL9.2), with a C-terminally fused foldon trimerization domain **(Fig 2a)**. Given the instability and poor recombinant expression yield observed for this ectodomain trimer construct (GPΔMuc), we opted to introduce two prefusion-stabilizing mutations previously described for EBOV and SUDV GPs (L579P and K588F)^37^ and one prefusion-stabilizing mutation that is broadly portable to all ebolavirus GPs (W615L)^38^. We thus compared the aforementioned construct to three designed GP mutant constructs harboring the L579P/W615L (termed BDBV GPΔMuc WL^2^P^4^), L579P/K588F (termed BDBV GPΔMuc F-P^4^), or L579P/K588F/W615L (termed BDBV GPΔMuc F-WL^2^P^4^) residue substitutions with or without co-transfection of furin. GP constructs co-expressed with furin led to complete cleavage of the GP into GP1 and GP2, while those expressed without furin resulted in incomplete GP cleavage **(Fig 2b)**. In presence of furin, all three stabilized BDBV GP constructs expressed to markedly greater levels than the unmutated construct with BDBV GPΔMuc F-WL^2^P^4^ yielding the greatest expression yields and monodispersity, as analyzed by size-exclusion chromatography and electron microscopy imaging of negatively stained samples **(Fig 2c-d)**. All ectodomains bound to immobilized EBOV-515 IgG **(Fig 2e)**, confirming the GPs are antigenetically intact, although the lower binding response observed for BDBV GPΔMuc, relative to the other three constructs, concurred with the heterogeneity observed by EM imaging (**Fig 2d-e**). These data enabled identification of an optimal prefusion-stabilization strategy for 2026 BDBV GP which will facilitate functional and immunological studies of this circulating virus.

**Figure 2.**
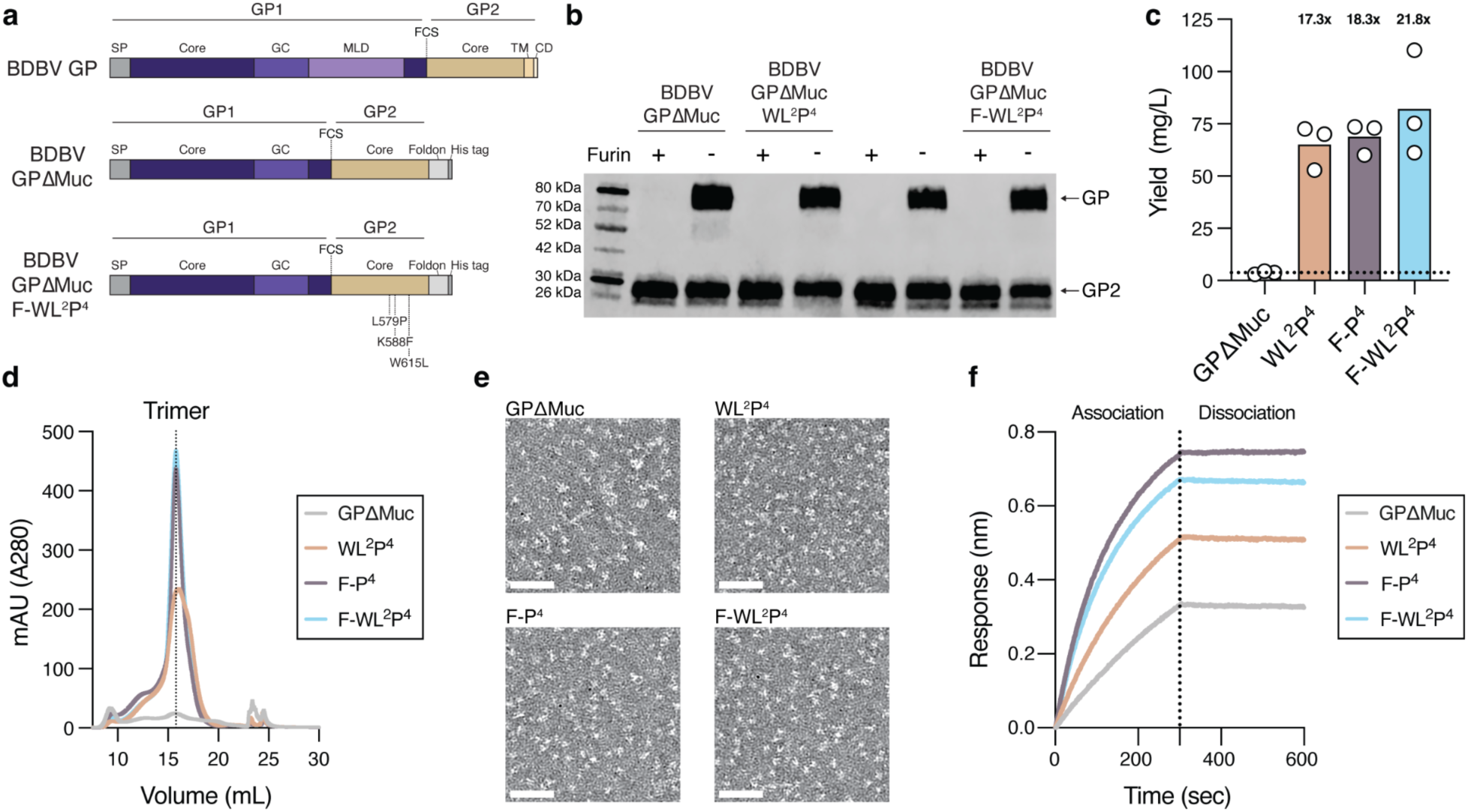
Design of a prefusion-stabilized BVDV GP ectodomain trimer. **a**, Schematic of BDBV GP domain organization and mutations included in GPΔMuc ectodomain constructs. CD, cytoplasmic domain; FCS, furin cleavage site; GC, glycan cap; MLD, mucin-like domain; TM, transmembrane domain. SP, signal peptide. The designed BDBV GPΔMuc constructs contain the following stabilizing mutations: WL^2^P^4^: L579P/W615L; F-P^4^: L579P/K588F; F-WL^2^P^4^: L579P/W615L/K588F. **b,** Western blot analysis of GP cleavage for the four designed BDBV GPΔMuc constructs using an anti-his tag primary antibody. **c,** Recombinant production yields for the BDBV GPΔMuc constructs after purification by affinity chromatography and size-exclusion chromatography. Each point represents an independently expressed biological replicate and the bar indicates the mean yield across the biological replicates. Fold-change in yields relative to BDBV GPΔMuc are indicated above each bar. **d,** Size-exclusion chromatograms for the BDBV GPΔMuc constructs. **e,** 30 Å low-pass filtered electron micrographs of negatively stained BDBV GPΔMuc ectodomains. Scale bar, 50 nm. **f,** Analysis of BDBV GPΔMuc constructs at a concentration of 0.03 mg/mL binding to immobilized EBOV-515 IgG assessed by BLI.

### Architecture of the prefusion BDBV GP trimer

We used single particle cryo-electron microscopy to determine a structure of BDBV GPΔMuc F-WL^2^P^4^ in complex with the EBOV-515 antibody Fab fragment at 3.2 Å resolution (**Fig 3a-b, Extended Data Fig 1 and Extended Data Table 1**). We selected EBOV-515 as this GP base-directed antibody is endowed with broadly neutralizing activity against EBOV, SUDV and historical BDBV isolates. Furthermore, EBOV-515 was shown to synergize with glycan cap-directed antibodies and to protect animals challenged with all three ebolavirus species prominently causing disease in humans^18,39^. The structure shows that GP adopts the prefusion state^22,23^, validating the design strategy, and resolves GP1, including part of the glycan cap, GP2 up to residue G598, and three bound EBOV-515 Fab fragments. Two out of the three introduced prefusion-stabilizing mutations are resolved in the cryoEM map, enabling visualization of their contribution in the context of BDBV GP. L579P introduces a proline residue that disfavors HR1 refolding to form a central 3-helix bundle (characteristic of postfusion GP) whereas K588F strengthens interactions with the GP1 N-terminal region, ultimately promoting prefusion BDBV GP stability **(Fig 3c)**.

**Figure 3.**
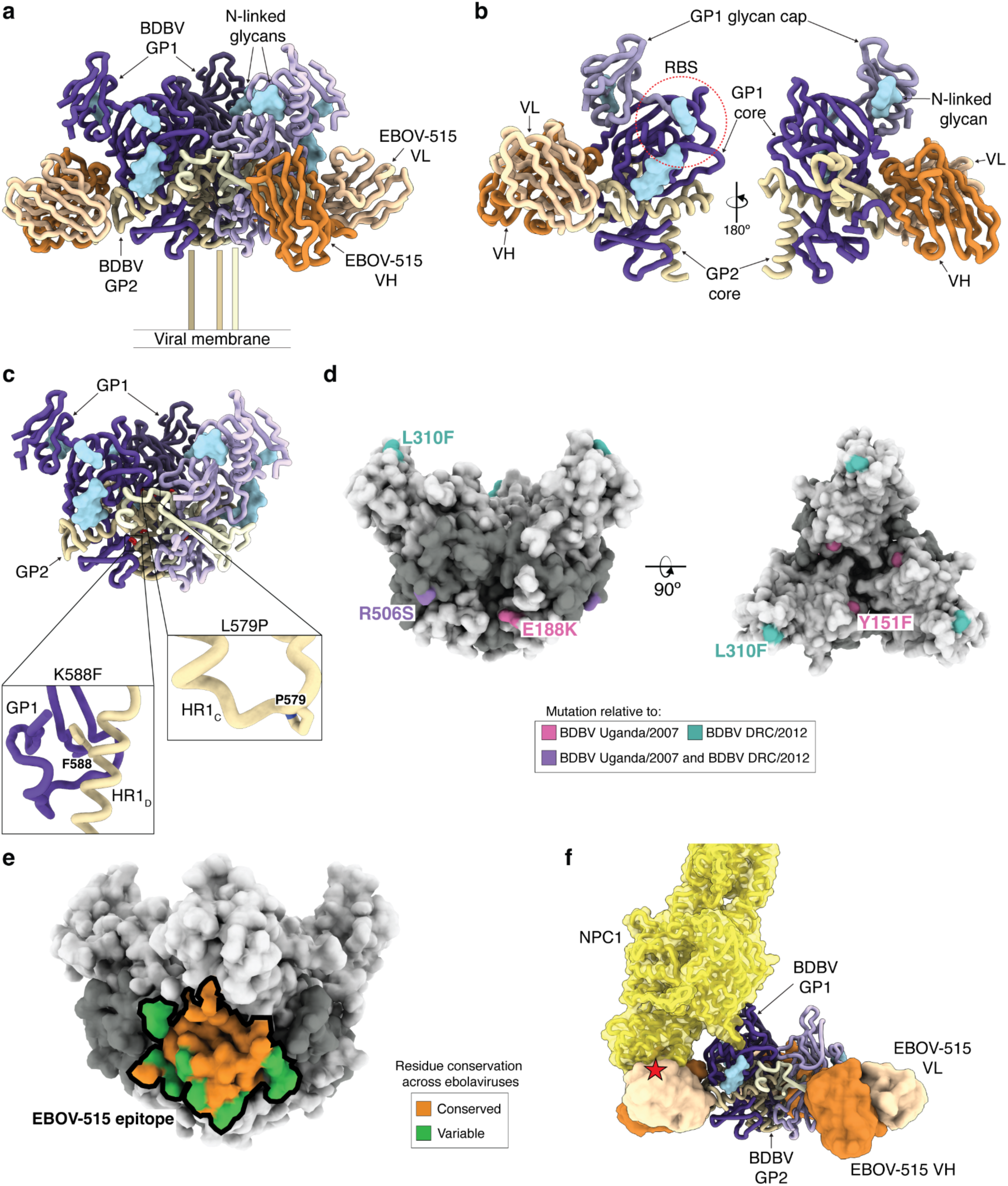
Architecture of the prefusion 2026 BDBV GP trimer bound to EBOV-515. **a**, CryoEM structure of the prefusion-stabilized (L579P/K588F/W615L) BDBV ectodomain trimer (<u>PP_006XHL9.2</u>) bound to three EBOV-515 Fab fragments. GP1 and GP2 are shown in shades of purple and gold, respectively. The EBOV-515 heavy chain is shown in orange and the light chain is shown in light orange (only the Fab variable domains were modelled into the density). N-linked glycans are rendered as light blue surfaces. The GP heptad repeat 2 and membrane-proximal region are not resolved in the cryoEM map and are schematically shown connecting to the viral membrane. **b,** A single BDBV GP protomer is shown bound to EBOV-515 in two distinct orientations with domains annotated. **c,** The BDBV ectodomain trimer is shown with zoomed-in views of the two prefusion-stabilized mutations resolved in the cryoEM density (L579P and K588F). **d,** Mapping of the 2026 lineage-defining GP mutations relative to 2007/2012 BDBV GPs on the cryoEM structure shown as a surface (GP1: light gray, GP2: dark grey) in two orthogonal orientations. **e,** Analysis of amino acid residue conservation within the EBOV-515 epitope among ebolavirus GPs colored according to the key (conserved: orange, variable: green) and mapped on the BDBV GP surface. **f,** Binding modes of the NPC1 receptor (yellow) and EBOV-515 (orange/light orange) to the BDBV GP. The position of NPC1 was determined by superimposing the EBOV GP–NPC1 (PBD: <u>5JNX</u>) and BDBV-EBOV-515 Fab structures. The EBOV GP trimer and the region of the BDBV GP glycan cap resolved in our structure (residues 191-310) are not shown for clarity. The red star denotes steric clashes.

The structure further resolves GP mutations distinguishing BDBV 2026 isolates from 2012 isolates (L310F and R506S) and from 2007 isolates (Y151F, E188K and R506S) **(Fig 3d)**. Residue 506 maps to the GP2 N-terminal region, and participates in antigenic sites targeted by several monoclonal antibodies, such as EBOV-515, EBOV-520 and ADI-15946 (recognizing the base of the fusion loop) as well as ADI-15878 and REGN-3479 (binding to the tip of the fusion loop). The Y151F GP1 residue substitution maps to the GP1 head and is solvent-exposed without obvious predicted impact on function, concurring with its occurrence in BDBV 2012 and EBOV GPs. The BDBV L310F mutation is in the glycan cap region that is targeted by a set of antibodies that use their CDRH2 or CDRH3 to displace the segment harboring residue 310, including BDBV289 (PDB 7KEJ) and BDBV-43 (PDB 7KEW)^39^. It is likely that the BDBV E188K mutation has a similar impact to the Q188R mutation, which was previously described to reduce binding of the BDBV91 mAb to EBOV soluble GP (sGP)^25^. sGP accounts for ∼80% of GP produced upon infection, forms homodimers and is believed to act as a decoy for antibodies^40–42^. However, BDBV91 does not bind to the prefusion GP trimer and is therefore non-neutralizing. Only three additional mutations (K299R, P344S, and I486T) have arisen in more than 10 of the 810 BDBV genomes sequenced in the current outbreak. K299R is in the glycan cap, but not resolved in our structure, P344S is in the mucin-like domain which was not included in the construct used here (given that it is unresolved in experimental structures), and I486T is in the region of GP1 preceding the furin cleavage, which is not resolved in our structure.

Our structural data reveal the molecular basis for EBOV-515 mediated broad neutralization of BDBV along with shared and unique features relative to complexes with EBOV GP, which shares ∼65% amino acid sequence identity with BDBV GP. EBOV-515 recognizes a glycan-free epitope involving contacts with both GP1 and GP2, including the base of the fusion loop **(Fig 3e)**. Most of the epitope residues are conserved across GPs from distinct ebolavirus species **(Fig 3e)**, explaining its broadly neutralizing activity of EBOV, SUDV and BDBV. S506 makes tenuous contacts with EBOV-515 in our structure **(Fig 3d)**, as is possibly the case for residue N506 in an EBOV-515-bound EBOV GP structure (PDB 7M8L)^18^. Moreover, EBOV GP N506 was modeled in the vicinity of the ADI-15946 heavy chain (PDB 6MAM)^43^ and of the REGN-3479 light chain whereas it does not appear to contact EBOV-520 (PDB 6OZ9)^44^ (PDB7TN9)^24^ in the respective complex structures. Therefore, our data suggest that amino acid mutations at position 506, such as the 2026 BDBV GP R506S substitution, may modulate binding and neutralization of some monoclonal antibodies recognizing this antigenic site in addition to modulating entry into cells.

Besides trapping GP in the prefusion state and reducing GP processing, as previously reported^17^, our data indicate that EBOV-515 would inhibit viral entry via steric hindrance with the NPC1 N-terminal domain **(Fig 3f)**, as previously described for the broadly neutralizing marburgvirus antibody MARV16 ^45^.

### Monoclonal antibody-mediated BDBV neutralization

To determine if monoclonal antibodies differentially recognize ebolavirus GPs, we assessed binding of a panel of immobilized antibodies to the BDBV DRC/2026 GPΔMuc F-WL^2^P^4^, EBOV Makona/2014 GPΔMuc, and SUDV Gulu/2000 GPΔMuc ectodomains by surface plasmon resonance **(Figure 4)**. Antibodies targeting the GP2 base and GP2 stalk exhibited comparable apparent binding affinities across the three GP ectodomains. Only 2 out of 6 antibodies targeting the GP1 glycan cap bound similarly to all three GPΔMuc ectodomains whereas the other 4 antibodies had marked variability across ebolavirus species. These results indicate that glycan cap antibodies are more sensitive to GP sequence variation, concurring with the more rapid evolutionary divergence of the glycan cap and mucin-like domain relative to other parts of GP. Comparison of cross-reactivity with the the wildtype BDBV DRC/2026 GPΔMuc F-WL^2^P^4^ ectodomain and the S506R BDBV DRC/2026 GPΔMuc F-WL^2^P^4^ ectodomain showed comparable binding for all antibodies tested despite the fact that residue 506 maps within the epitope of several GP2 base antibodies **(Figure 4)**. We further found that GP2 base and stalk-directed antibodies recognized the BDBV DRC/2026 GPΔMuc F-WL^2^P^4^ and SUDV Gulu/2000 GPΔMuc ectodomains cleaved by thermolysin, to remove the GP1 glycan cap as mediated by endosomal cathepsins during viral entry. However, no binding of glycan cap-directed antibodies was observed, as expected due to shedding of this domain upon cleavage. Three of the GP2 base antibodies, ADI-23744, ADI-15946, and EBOV-520, displayed markedly increased affinities towards the thermolysin-cleaved GP ectodomains compared to the uncleaved ectodomains, consistent with the glycan cap partially shielding the epitopes of these antibodies. Overall, our data show that numerous neutralizing antibodies retain strong binding to the currently circulating BDBV DRC/2026 GP.

**Figure 4.**
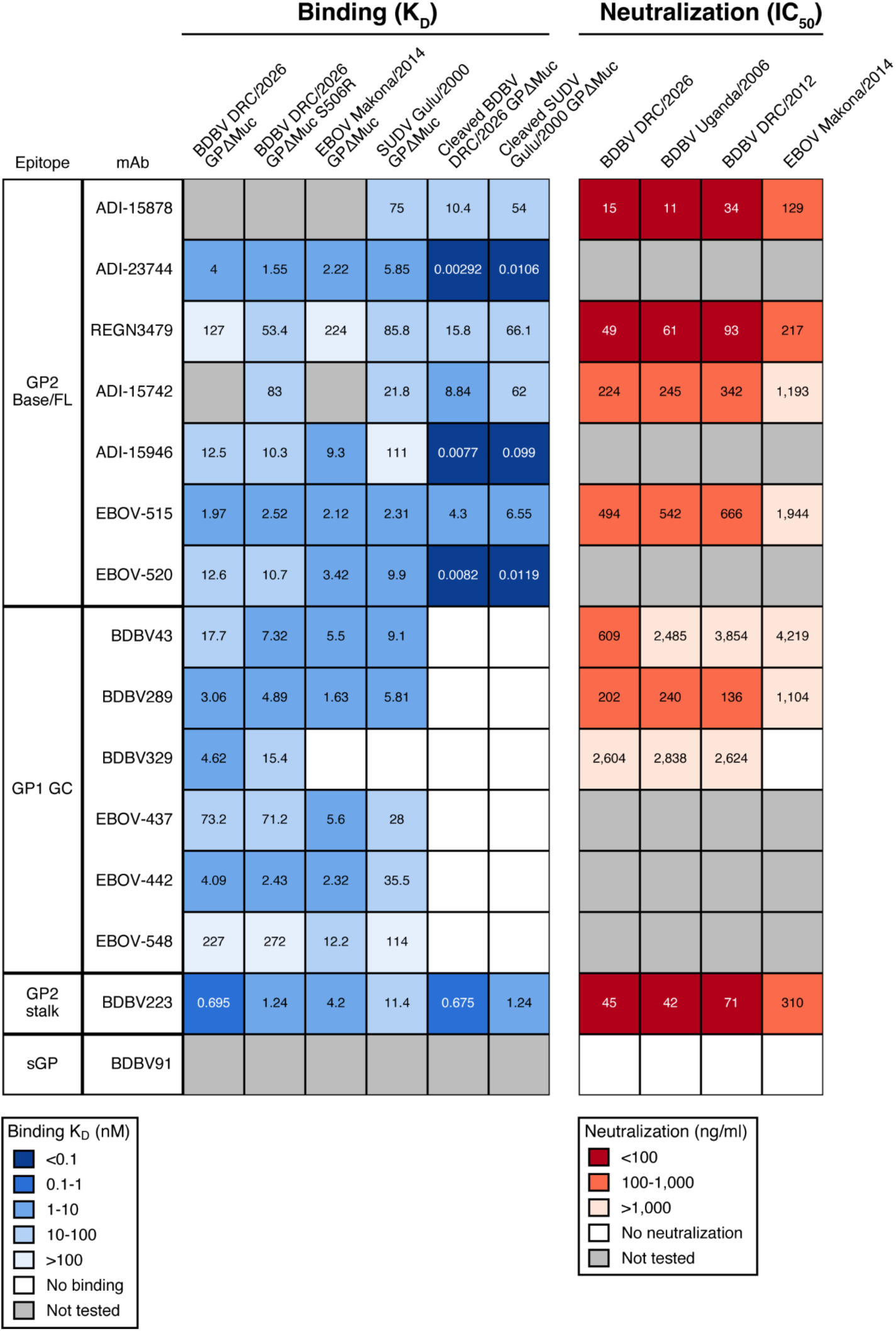
Evaluation of the BDBV 2026 antigenicity. Apparent binding affinity and neutralization potency of monoclonal antibodies targeting ebolavirus GPs. Binding affinity was assessed by surface plasma resonance using the BDBV DRC/2026 GPΔMuc F-WL^2^P^4^, BDBV DRC/2026 GPΔMuc F-WL^2^P^4^ harboring the S506R mutation, EBOV Makona/2014 GPΔMuc, SUDV Gulu/2000 GPΔMuc, thermolysin cleaved BDBV DRC/2026 GPΔMuc F-WL^2^P^4^, or thermolysin cleaved SUDV Gulu/2000 GPΔMuc ectodomain. One to two technical replicates were performed to determine the binding affinity (K_D_; nM) Neutralization assays were performed using pseudoviruses harboring the GP proteins of BDBV DRC/2026, BDBV Uganda/2007, BDBV DRC/2012, and EBOV Makona/2014. The assay was performed in quadruplicate for each antibody at each concentration to determine the 50% inhibitory concentration (IC50; ng/ml), and the assay was independently repeated three times. The average of IC50 values from three independent assays is shown. FL: fusion loop; GC: glycan cap; sGP: soluble glycoprotein.

To assess whether sequence variation in the 2026 BDBV GP alters sensitivity to monoclonal neutralizing antibodies, we compared neutralization of contemporary and historical BDBV GP pseudoviruses using a lentiviral pseusotyping platform (**Table 1 and Extended Data Fig 2**). We observed that the BDBV DRC/2026 isolate remained susceptible to all antibodies that neutralized earlier BDBV isolates. Among the therapeutic antibodies tested, both ADI-15878 (a component of MBP134) and REGN3479 (also known as maftivimab and a component of Inmazeb) potently neutralized BDBV DRC/2026 pseudovirus. These findings indicate that the 2026 BDBV GP retains susceptibility to the broadly neutralizing antibodies tested, including components of the clinical stage MBP134 cocktail and of the Inmazeb cocktail approved for EBOV.

## Discussion

Although the licensed Ervebo vaccine has proven highly effective during the pivotal Ebola Ça Suffit! Clinical trial in Guinea in 2015^29^, it is based on EBOV GP, which is considerably divergent from BDBV GP and even more so from SUDV GP. Consequently, the protective efficacy of this vaccine against disease caused by BDBV and SUDV in humans remains uncertain and the World Health Organization announced it will be supporting its clinical evaluation in the Democratic Republic of the Congo where the ongoing outbreak of BDBV-induced disease is spreading rapidly (https://www.who.int/news/item/20-08-2026-who-and-africa-cdc-welcome-the-allocation-of-ebola-vaccines-to-the-democratic-republic-of-the-congo). The marked genetic and antigenic distance between GP trimers of various ebolavirus species, however, indicates that species-matched vaccines will need to be developed for providing optimal protection against BDBV, as has been initiated for SUDV^46,47^.

Prefusion-stabilization has enabled recombinant production of ectodomain constructs faithfully recapitulating the native architecture and antigenicity of numerous viral surface glycoproteins. A combination of disulfide engineering and cavity filling mutations yielded several approved respiratory syncytial vaccines^48–53^ whereas introduction of proline residues incompatible with the large-scale fusogenic conformational changes led to several efficacious COVID-19 vaccines^54–59^. We identify here an optimal prefusion GP stabilization strategy by introducing mutations ported from other ebolavirus GPs to BDBV GP^37,38^. A minimum of two mutations were needed to enhance recombinant expression yields by greater than 10-fold, relative to an unmutated ectodomain construct. Specifically, substitution of GP2 residue 579 for a proline, to disfavor formation of the postfusion HR1 central 3-helix bundle, and of GP2 residue 588 for a phenylalanine, to strengthen interactions with GP1, acted in concert to promote prefusion BDBV GP stability. We anticipate that the stabilized BDBV GP ectodomain trimer described here will accelerate the design of BDBV vaccines and immunological studies of infected or vaccinated individuals or animals to assist with the identification of potent BDBV GP monoclonal antibody therapeutics.

Naturally occurring GP mutations have been documented by sequencing isolates found in humans infected with the various ebolavirus species associated with epidemics^60,61^. The glycan cap and the mucin-like domain are prime targets of serum antibodies and the vast majority of GP mutations accumulate in these domains, as is the case with recently sequenced 2026 BDBV genomes. Mutations in other GP regions, such as the GP1 head or the GP1/GP2 base have also been observed and it is therefore key to map these mutations and characterize them experimentally. The data presented here provide a molecular blueprint and a functional framework to understand present and future GP mutations and their impact on immune responses and pathogenicity.

The development of mAbs that are neutralizing and protective against other ebolavirus species than EBOV, particularly BDBV and SUDV, inclusive of all their variants, is a priority to identify therapeutic solutions for the repeated spillovers of these latter two viruses. The identification and preclinical evaluation of several mAbs with broad activity against EBOV, BDBV and SUDV, based on direct viral neutralization and Fc-mediated effector functions, is a key step forward toward achieving this goal^14–18,20,22–24,26^. Several mAbs targeting the GP base are endowed with potent and broad neutralizing activity and provide effective post-exposure protection in animal models whereas many mAbs targeting the glycan cap have limited *in vitro*neutralizing activity and modest therapeutic efficacy although they can synergize with other mAbs. The most advanced candidate is a broadly protective mAb cocktail (designated MBP134) which comprises derivatives of ADI-15878 and ADI-15946. Several other mAb cocktails have yielded promising results for post-exposure treatment of animals experimentally infected with EBOV, BDBV and SUDV and warrant clinical evaluation. The observed binding and *in vitro* neutralizing activity against 2026 BDBV for the panel of mAbs evaluated here provides encouraging results as to the expected retention of *in vivo* efficacy of these therapeutic candidates. Specifically, the equivalent 2026 BDBV pseudovirus neutralization potency of of ADI-15878 to that observed against prior BDBV isolates and 2014 EBOV provides support to the recent decision of the World Health Organization to evaluate if administration of MBP134 and remdesivir improves survival among patients diagnosed with BDBV infection in the Democratic Republic of the Congo (https://www.who.int/news/item/02-07-2026-patient-enrolment-begins-in-a-scientific-trial-to-identify-the-first-effective-treatments-for-bundibugyo-virus-disease).

## Methods

### Epidemiological surveillance data

Weekly counts of confirmed BDBV disease cases and deaths were obtained from the Weekly External Situation Reports on the Bundibugyo virus disease outbreak, published by the World Health Organization Regional Office for Africa (WHO AFRO: https://www.afro.who.int). Cumulative numbers of confirmed cases and confirmed deaths were manually extracted from Situation Reports 01–18. Suspected and probable cases were excluded. Data were visualized in R v.4.4.1 using the ggplot2 package v.4.0.1.

### Phylogenetic and sequence analysis

BDBV sequence data were downloaded from Pathoplexus (https://pathoplexus.org) on 28 August 2026. All entries released under the OPEN data use terms were retrieved, together with one sequence released under RESTRICTED terms (Pathoplexus accession no. PP_006XXY5.1), which was included with permission from the data submitters (Prof. Dr. Christian, Charité – Universitätsmedizin Berlin, Germany). In total, n = 46 sequences were analyzed. Sequences were aligned with MAFFT v7.526^62^ using the ––localpair option. Maximum-likelihood phylogenies were reconstructed from the whole-genome and GP coding region nucleotide alignments using IQ-TREE v3.0.1^63^. For each alignment, the best-fitting nucleotide substitution model was selected with ModelFinder as implemented in IQ-TREE based on the Bayesian information criterion (BIC), with FreeRate models included to account for among-site rate heterogeneity (-m TEST-mrate R). TIM+F+R2 and TN+F+R2 were selected for the whole-genome and GP trees, respectively. Branch support was assessed with 1,000 ultrafast bootstrap replicates. Trees were visualised in PearTree v1.2.0 (https://github.com/artic-network/peartree), and amino acid substitutions were plotted with snipit v1.7 (https://github.com/aineniamh/snipit).

### Cells

HEK293T cells (a human embryonic kidney cell line; ATCC, Cat# CRL-1573) and Lenti-X 293T cells (Takara, Cat# 632180) were cultured in high glucose Dulbecco’s Modified Eagle Medium (DMEM) (Wako, Cat# 044-29765) supplemented with 10% fetal bovine serum (FBS) (Nichirei, Cat# 175012) and 1% penicillin/streptomycin (P/S) All cells were maintained at 37°C in a humidified atmosphere containing 5% CO_2_. Expi293 cells (ThermoFisher) were grown in Expi293 medium at 37°C at 5% CO_2_. Cell lines were not authenticated or tested for mycoplasma contamination.

### Constructs

The construct encoding the wild-type BDBV GPΔMuc ectodomain (residues 1-312 and 471-649) with a C-terminal T4 foldon trimerization sequence followed by an 8x His tag was codon-optimized, synthesized, and inserted into pcDNA3.4 by Genscript. The stabilized constructs containing the L579P and K588F, L579P and W615L, or L579P, K588F, and W615L mutations were generated through mutagenesis of the wild-type BDBV GPΔMuc by Genscript. The construct encoding the EBOV Makona/2014 GPΔMuc, SUDV Gulu/2000 GPΔMuc, furin, the EBOV-515 heavy chain, and the EBOV-515 light chain was previously described^45^.

A plasmid expressing the Ebola virus GP protein of the Makona strain (EBOV Makona/2014) was prepared in the previous study^64^. Plasmids expressing the BDBV GP proteins of the Uganda/2007 strain (Accession no. FJ217161.1), DRC/2012 strain (Accession no. MT680262.1), and DRC/2026 strain (Pathoplexus accessions no. PP_006XXY5.1) were artificially synthesized by the Fasmac Gene Synthesis service (Fasmac Co., Ltd.). Permission to use the 2026 GP sequence in this study was obtained from the data submitters, Prof. Dr. Christian Drosten (Charité – Universitätsmedizin Berlin, Germany) Plasmids expressing the BDBV GP proteins of the 2026 strain derivative were generated by site-directed overlap extension PCR using the primers listed in **Extended Data Table 2**. The resulting PCR fragment was digested with KpnI and NotI and inserted into the corresponding site of the pCAGGS vector^65^.

For monoclonal antibodies used in the neutralization assays, plasmids expressing the heavy chain (HC), kappa chain (KC), or lambda chain (LC) were constructed as follows. The sequences of the variable regions were obtained from the RCSB Protein Data Bank (https://www.rcsb.org/) for REGN-3479 (PDB: 7TN9), BDBV43 (PDB: 7KEW), and BDBV91 (PDB: 5KEM); from IEDB

(https://www.iedb.org/) for ADI-15742 and EBOV-515; and from EBOV-AbDab (https://zenodo.org/records/10564972) for BDBV223, BDBV289, and BDBV329. The variable regions of REGN-3479 were generated by overlap PCR using the primers listed in **Extended Data Table 2**. The variable regions of the HC and KC of the other antibodies, and the open reading frame of the BDBV289 LC, were artificially synthesized together with the corresponding leader sequences (MEFGLSWLFLVAILTGVQC for HC and MDMRVPAQLLGLLLLWFPGARC for KC) (Fasmac). The resulting coding sequences of the variable regions of the heavy and light chains were cloned into the pCAGGS vector together^65^ with the sequence encoding the human immunoglobulin γ1 constant region (Accession no. QRG33624.1) or the kappa constant region (Accession no. AAA59000.1). Nucleotide sequences were determined by DNA sequencing services (Eurofins Genomics), and the sequence data were analyzed by SnapGene v8.2.2 software (MDF Co., Ltd.). For monoclonal antibodies used for surface plasmon resonance, the variable regions of heavy and light chain sequences were synthesized and inserted into the pTT5 vector containing the constant regions for the respective heavy, lambda, or kappa chain by Genscript.

### Recombinant protein expression and purification

The recombinant BDBV, EBOV, and SUDV GPΔMuc ectodomains were expressed and purified as previously described^45^. In brief, Expi293 cells were grown to 3 million cells per mL and transfected with the BDBV GPΔMuc construct with or without furin (at a 3:1 mass ratio, if co-transfected furin) using Expifectamine293 (ThermoFisher) according to the manufacturer’s instructions. After 5 days, the supernatant was collected, clarified by centrifugation, and incubated with Ni Sepharose Excel Resin (Cytiva) for 1 hour at room temperature. The resin was collected in a gravity-flow column and washed with wash buffer (25 mM Na_2_PO_4_, 300 mM NaCl, 50 mM imidazole, pH 8.0). The GPΔMuc was eluted from the resin using elution buffer (25 mM Na_2_PO_4_, 300 mM NaCl, 500 mM imidazole, pH 8.0) and further purified and exchanged in TBS (20 mM Tris-HCl, 100 mM NaCl, pH 7.4) using a Superose6 Increase 10/300 GL column. The purified proteins were flash frozen and stored at –80°C until use.

Thermolysin cleavage of the BDBV and SUDV GPΔMuc ectodomains was completed as previously described^45^. In brief, the ectodomains were expressed and purified as described above. The GPΔMuc ectodomains were diluted to 1 mg/mL in TBS and incubated thermolysin (Sigma-Aldrich) diluted to 0.2 mg/mL at 37°C for 1 hr. Next, phosphoramidon (Sigma-Aldrich) was added to a final concentration of 500 µM to inhibit the thermolysin. The cleaved GPΔMuc ectodomains were purified using a Superdex 200 Increase 10/300 GL column into TBS.

EBOV-515 and ADI-15878 IgGs were expressed and purified as previously described^45^. Briefly, Expi293 cells at a density of 3 million cells per mL were co-transfected with the EBOV-515 heavy and light chain constructs at a 1:1 mass ratio using Expifectamine293 (ThermoFisher) according to the manufacturer’s instructions. After 4 days, the supernatant was collected, clarified by centrifugation, and flowed over a HiTrap Protein A column (Cytiva) for 1 hour at room temperature. The column was washed with 20 mM Na_2_PO, pH 8.0 and the recombinant IgG was eluted using 0.1 M citric acid, pH 3.0 directly into 1 M Tris-HCl, pH 9.0. The IgG was exchanged into TBS (20 mM Tris-HCl, 150 mM NaCl, pH 8.0) using a 100-kDa Amicon Ultra centrifugal filter. To generate EBOV-515 Fab, the IgG was digested with LysC (NEB), resuspended in TBS, at a 4,000:1 mass ratio at 37°C for 18 hours. Undigested IgG and free Fc fragments were collected by incubating the digestion product with MabSelect resin (Cytiva) for 1 hour at room temperature. The resin was captured with a gravity-flow column and the flow-through was collected. To further purify the Fab, the flow-through was run over a Superdex75 Increase 10/300 GL column equilibrated with TBS. The IgG and Fab were stored at 4°C until use.

The additional monoclonal antibodies used for surface plasmon resonance were expressed and purified from CHO-K1 cells into a sodium acetate buffer with 0.2 M L-Arginine, pH5.5 by Genscript.

### Surface plasmon resonance (SPR)

To assess the binding affinity between the monoclonal antibodies and the GPΔMuc ectodomains, the monoclonal antibodies were diluted in HEPES buffered saline with Tween-20 (HBST; Carterra) to 1, 0.2, 0.04 µg/mL and spotted onto a pre-conditioned Protein A-coated biosensor chip (PAHC200M; Carterra). Next, a non-regenerative kinetics capture experiment was performed by injecting the GPΔMuc ectodomain serially diluted in HBST with 0.5% BSA from 0.411 to 900 nM over the surface of the mAb-coated biosensor chip with an association time of 300 s and a dissociation time of 900 s for each concentration of GPΔMuc ectodomain. All injections were completed at 25°C. The chip was regenerated with 10 mM glycine, pH 2.0 and re-spotted with monoclonal antibodies before proceeding with the next GPΔMuc ectodomain kinetics capture cycle. The resulting data was analyzed using the Kinetics software (Carterra). The data were corrected using a reference spot and leading buffer blank, y-aligned to the baseline before the first GPΔMuc ectodomain injection, and fit using a 1:1 binding model using float T_0_ to account for the non-regenerative kinetics cycle.

### Cryo-EM sample preparation and data collection

BDBV GPΔMuc-EBOV-515 Fab complexes were formed by incubating 100 µg of BDBV GPΔMuc with 150 µg of EBOV-515 at 4°C for 18 hours. Excess Fab was removed by washing the complexes 5 times with TBS in a 100-kDa Amicon Ultra centrifugal filter. Immediately prior to freezing, either CHAPSO at a final concentration of 6 mM was added to the GP-Fab complexes concentrated to 4 mg/mL or a surfactant mixture (0.0048% FOM, 0.06% Bril-35, 0.05% CHAPSO and 0.25% Amphipol 8-35)^66^ was added to the GP-Fab complexes concentrated to 1 mg/mL. The complexes were added to the freshly glow-discharged UltrAuFoil R 2.0/2.0 grids (200 mesh) and the grids were then plunge frozen using a Vitrobot MarkIV (ThermoFisher) with a 10 second wait time, a –2 blot force, and a 4 second blotting time at 100% humidity and 23°C. Data were acquired on an FEI Glacios operated at 100 kV and equipped with a Gatan K3 Summit. Automated data collection was carried out using a Leginon at a nominal magnification of 45,000x with a pixel size of 0.89 Å, a defocus range of –0.1 to –4.0 µm, and a stage tilt of 0° or 30°.

### Cryo-EM data processing

Patch motion correction. and estimation of microscope CTF parameters were completed using cryoSPARC (v5.0.6)^67^. Particle picking was carried out using 20 Å low-pass filtered templates generated from a prior EBOV GP-EBOV515 Fab model (PBD: 7KF9)^39^ followed by extraction of particles downsampled to a pixel size of 1.78 Å using a box size of 200 pixels^2^ were performed in cryoSPARC. Two rounds of reference-free 2D classification followed by ab-initio 3D and heterogenous refinement were performed to select well-defined particles using cryoSPARC. The pixel size was adjusted to 0.89 Å and box size was adjusted to 400 pixels, after which 3D refinement was carried out using non-uniform refinement with C3 symmetry and global and per-particle defocus in cryoSPARC^68^. Next, reference-based motion correction followed by another round of nonuniform refinement with C3 symmetry and global and per-particle defocus were performed with cryoSPARC. Local refinement with a mask encompassing the BDBV GP excluding the HR2 stalk (residues 612-649) and the variable regions of the EBOV-515 and enforcing C3 symmetry was conducted using cryoSPARC.

### Model building and refinement

A previously determined BDBV GP (PDB 7KEW)^39^ was docked into the cryoEM reconstruction using ChimeraX^69^ before manual rebuilding using Coot^70^ and refinement using Phenix^71^ using sharpened and unsharpened maps. Validation was carried out with Phenix, Molprobity^72^ and Privateer^73^.

### Monoclonal antibody preparation

Among the anti-EBOV monoclonal antibodies, ADI-15878 was purchased from MedChemExpress (Cat# HY-P991450), and the others were constructed in-house. Recombinant immunoglobulin G (IgG) was produced and purified as previously described^74^. Briefly, pairs of heavy– and light-chain plasmids were co-transfected into 293T cells using PEI MAX (Polysciences, Inc.). The culture medium was replaced with fresh D-MEM (High Glucose; Wako) the next day of the transfection, and the supernatant was harvested 3 days post-transfection. IgGs were purified using MonoSpin ProA Spin columns (GL Sciences).

### Neutralization assay

Pseudoviruses were prepared as previously described^75–79^ with minor modifications. Briefly, lentivirus (HIV-1)-based, luciferase-expressing reporter viruses were pseudotyped with the EBOV/BDBV GPs. LentiX-293T cells (1,000,000 cells) were cotransfected with 1 μg psPAX2-IN/HiBiT^80^, 1 μg pWPI-Luc2^80^, and 500 ng plasmids expressing parental GPs using TransIT-293 transfection reagent (Mirus, Cat# MIR2704) according to the manufacturer’s protocol. Two days post transfection, the culture supernatants were harvested and filtrated. The pseudoviruses were stored at –80°C until use. Neutralization assay was prepared as previously described^75–79^. Briefly, the EBOV/BDBV GPs pseudoviruses (counting ∼20,000 relative light units) were incubated with serially diluted monoclonal antibodies in a range from 50 µg/ml to 5.0 × 10-9 µg/ml at the final concentration at 37°C for 1 h. Pseudoviruses without sera were included as controls. Then, a mixture of pseudovirus and antibody was added to HEK293T cells (10,000 cells) in a 96-well white plate. At 2 d.p.i., the infected cells were lysed with a Bright-Glo™ Luciferase Assay System (Promega, Cat# E2650), and the luminescent signal was measured using a GloMax explorer multimode microplate reader 3500 (Promega). The assay of each serum sample was performed in quadruplicate, and the half maximal Inhibitory Concentration (IC50) was calculated using Prism 9 (GraphPad Software).

## Supplementary Figures

**Extended Data Figure 1.**
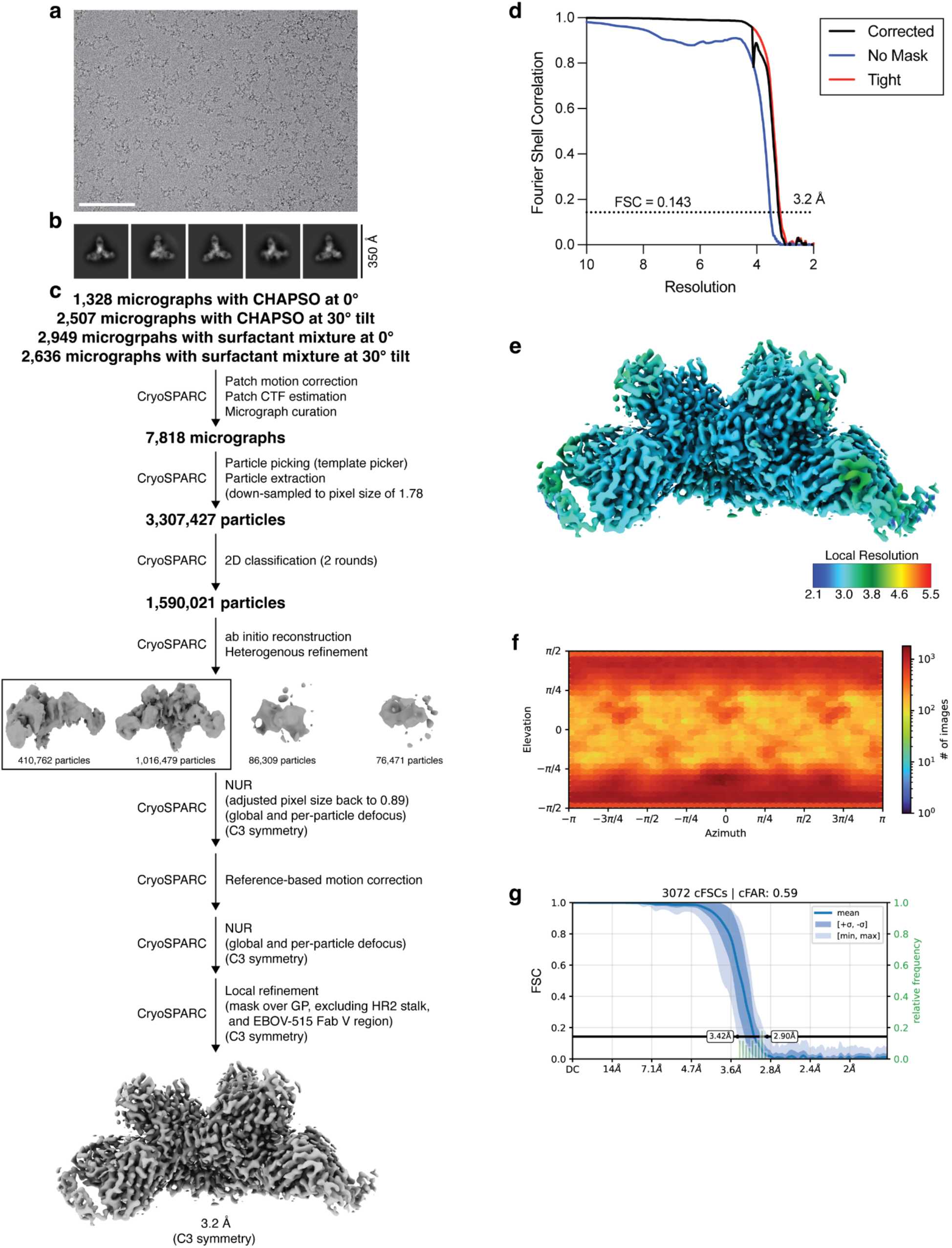
Cryo-EM data processing workflow for the BDBV GPΔMuc F-WL^2^P^4^-EBOV-515 Fab complex. **a-b**, Representative cryo-EM micrograph (a) and 2D class averages (b) for the BDBV GPΔMuc F-WL^2^P^4^-EBOV-515 Fab complex. Scale bar: 100 nm. **c,** Cryo-EM data processing workflow. NUR: non-uniform refinement. **d,** Gold-standard fourier shell correlation curves for the locally refined BDBV GPΔMuc F-WL^2^P^4^-EBOV-515 Fab complex Fab complex (using a mask comprising the GP trimer, excluding the HR2 stalk, and EBOV-515 Fab variable domains). **e,** Local resolution calculated with cryoSPARC for the locally refined BDBV GPΔMuc F-WL^2^P^4^-EBOV-515 complex. **f,** Heat map of angular distribution for the particles contributing to the final reconstruction. **g,** Conical fourier shell correlation plot.

**Extended Data Figure 2.**
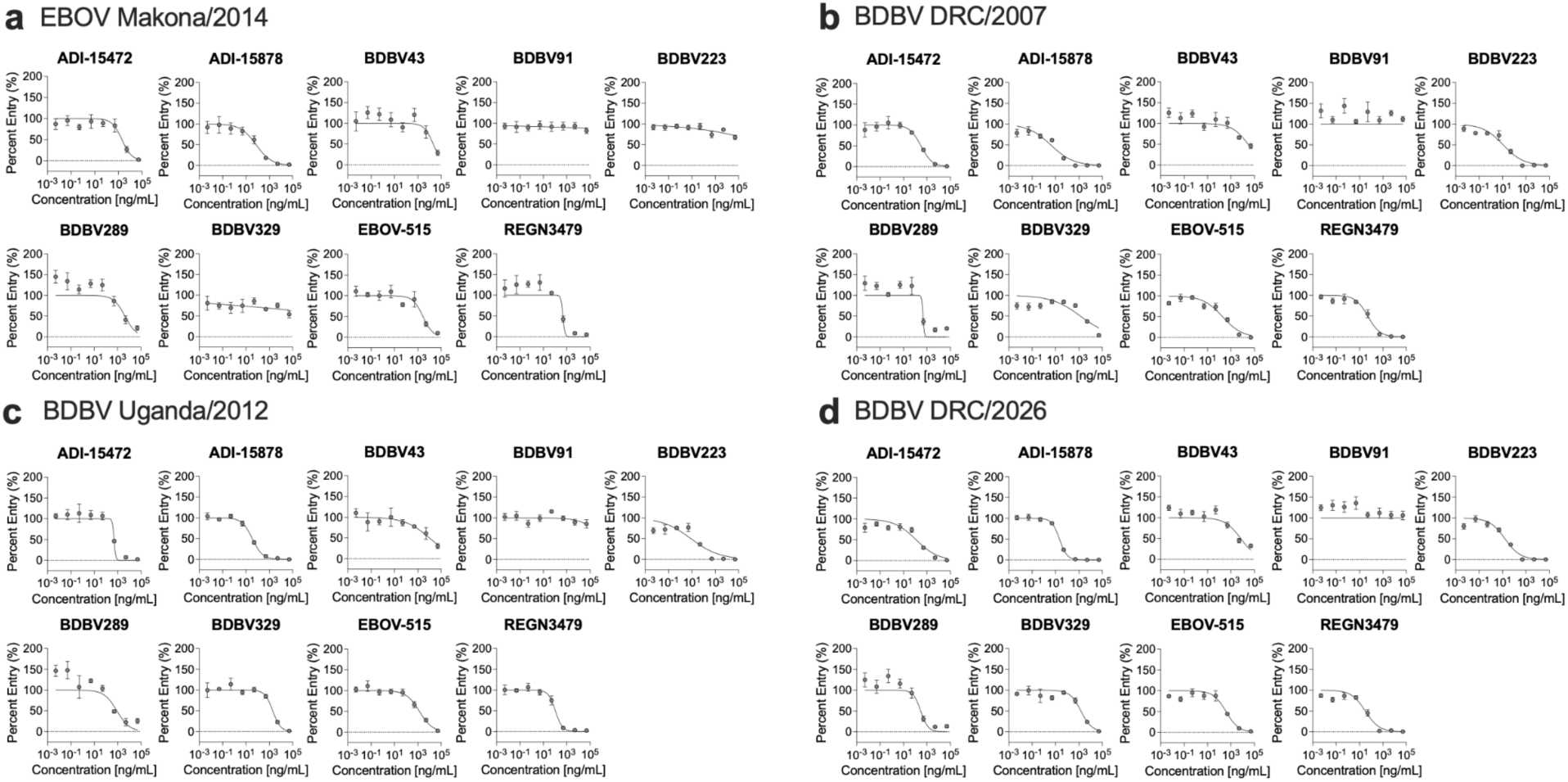
Neutralization dose-response curves for monoclonal antibodies. **a-d**, Representative neutralization dose-response curves for ADI-15472, ADI-15878, BDBV43, BDBV91, BDBV223, BDBV289, BDBV329, EBOV-515, and REGN3479 against HIV-based pseudovirus displaying the EBOV Makona/2014 (a), BDBV DRC/2007 (b), BDBV Uganda/2012 (c), or BDBV DRC/2026 (d) GP.

**Extended Data Table 1.**
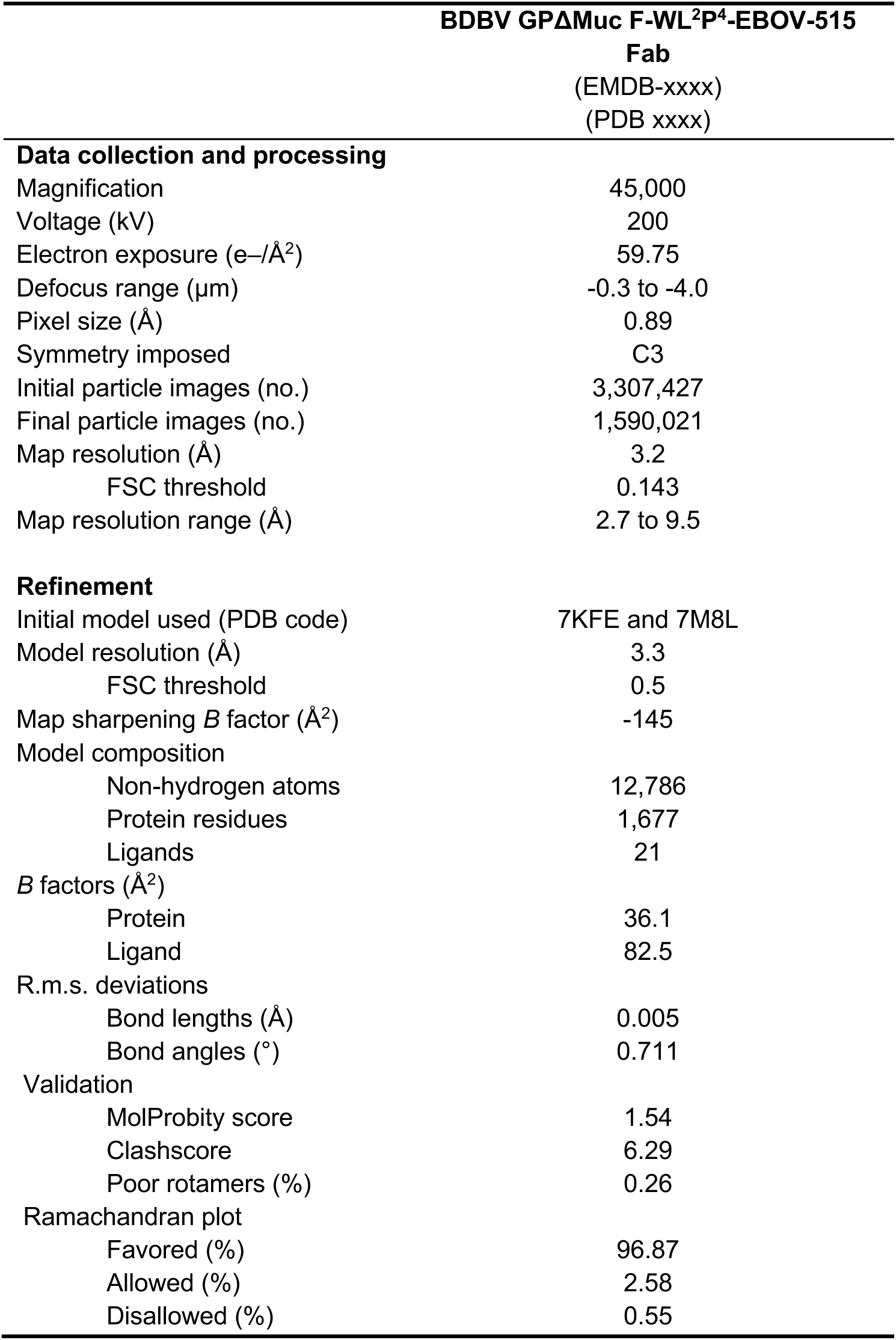
Cryo-EM data collection, refinement and validation statistics.

**Extended Data Table 2.**
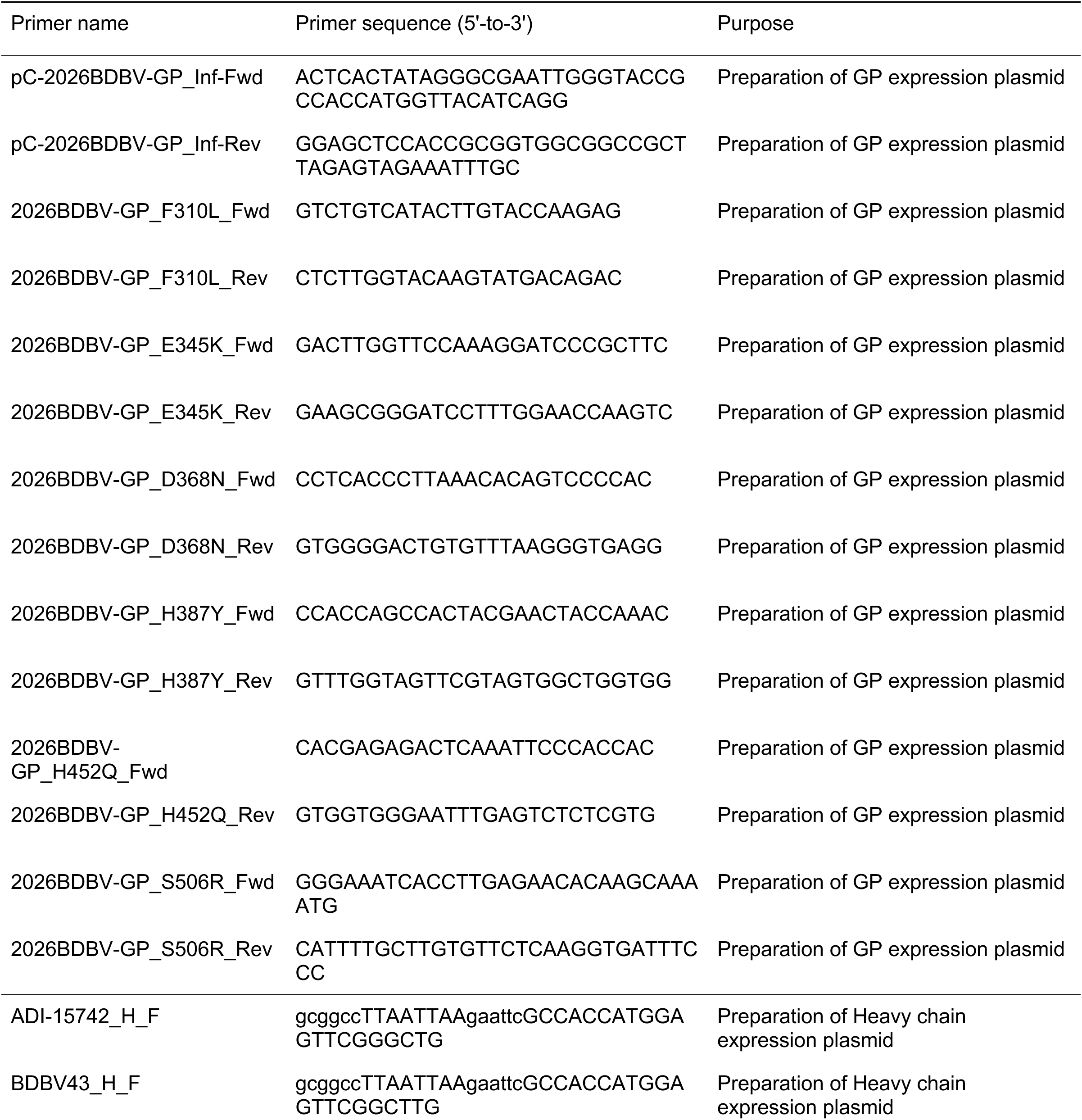
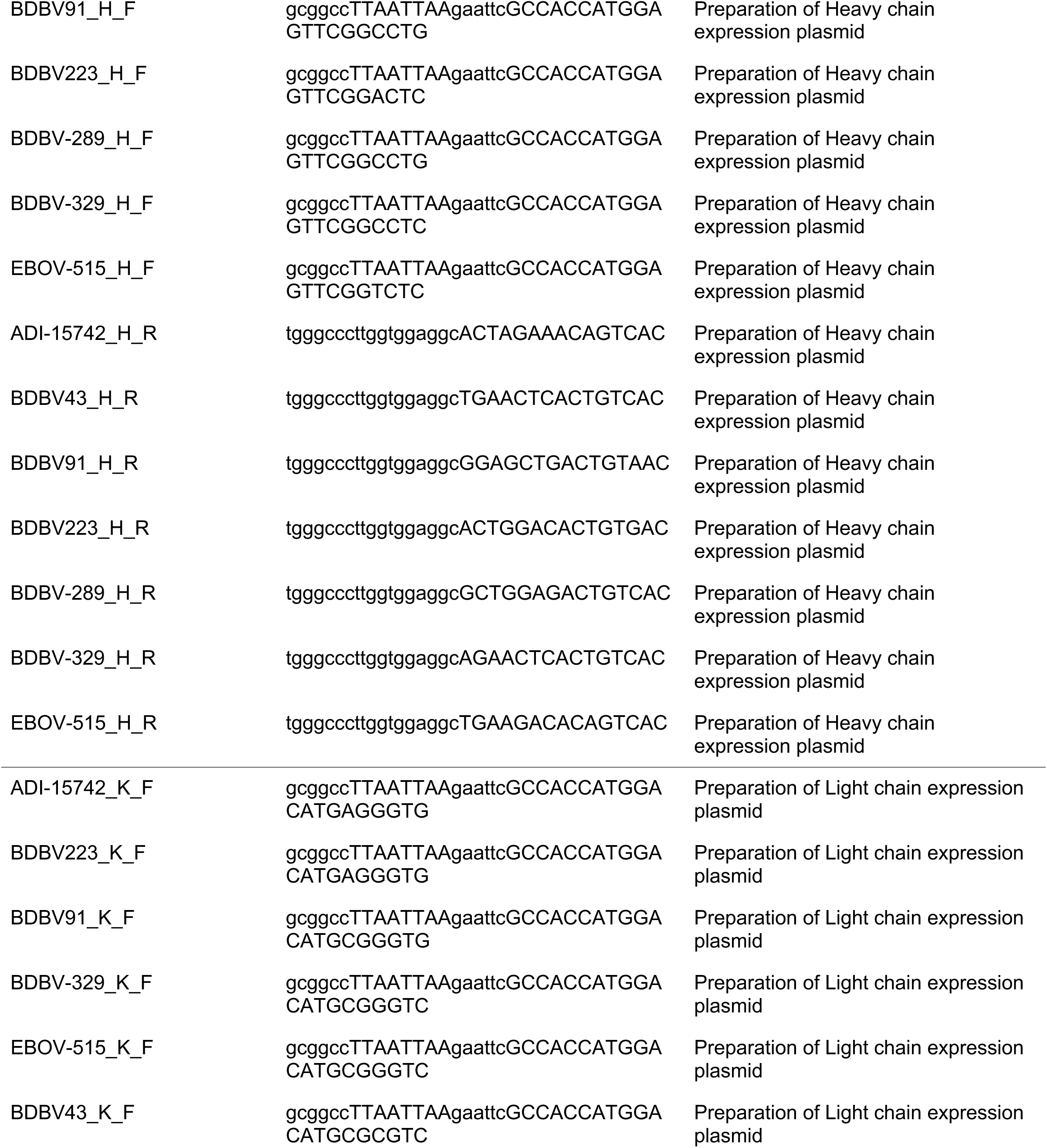
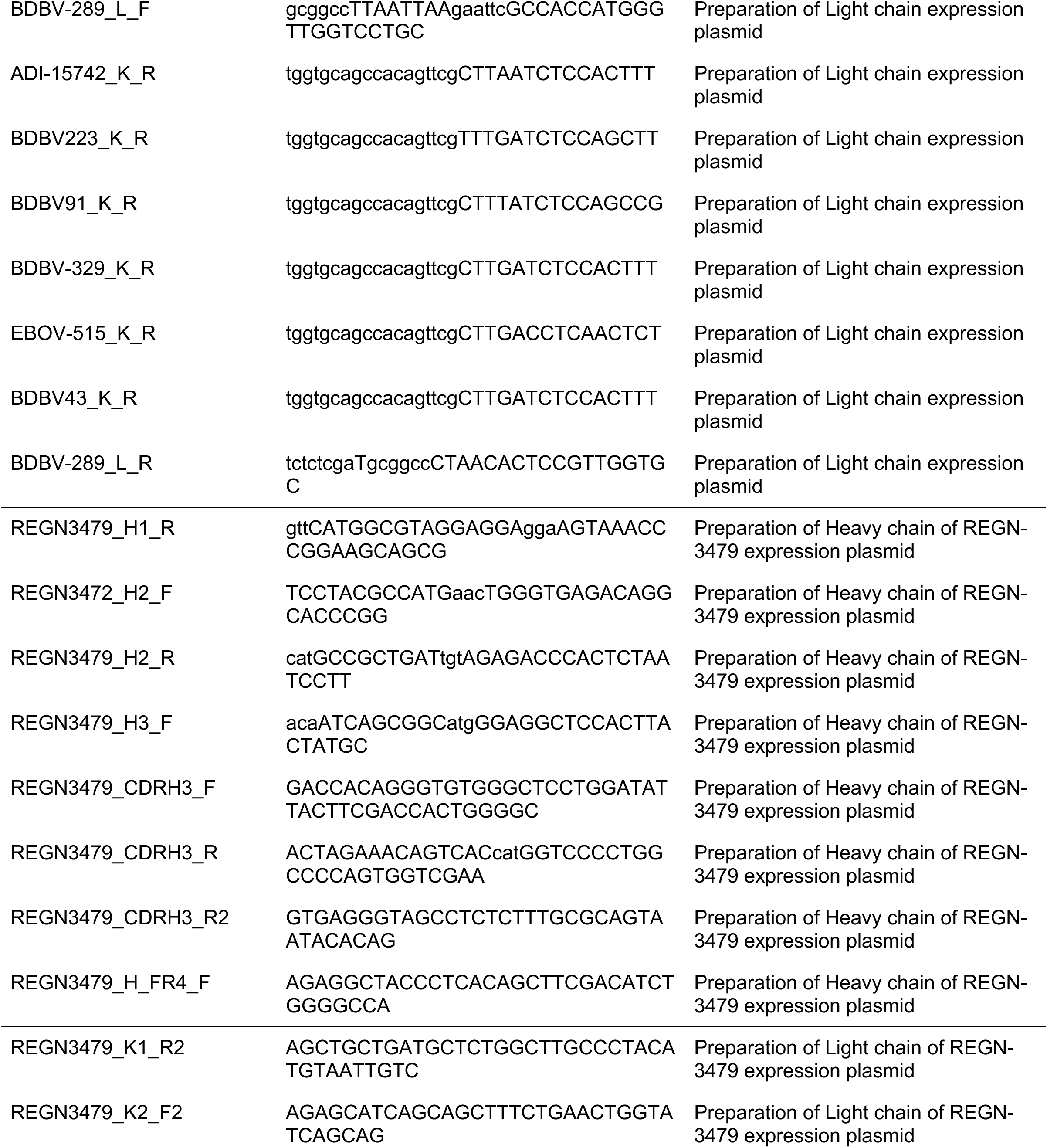
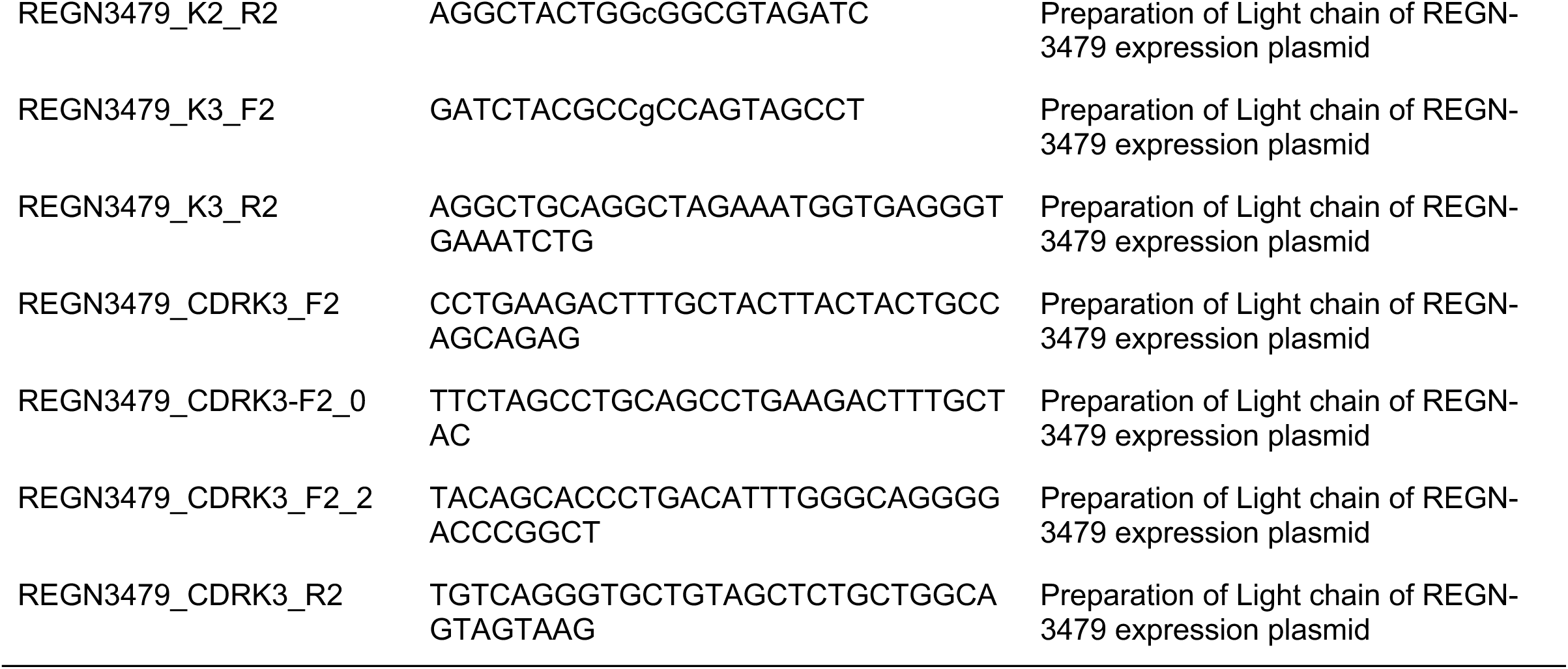
Primers used in this study.

